# Chronic deregulation of the spindle assembly checkpoint triggers myelosuppression and gastrointestinal atrophy

**DOI:** 10.1101/2023.09.08.556836

**Authors:** Gerlinde Karbon, Fabian Schuler, Manuel Haschka, Mathias Drach, Rocio Sotillo, Stephan Geley, Diana C.M. Spierings, Andrea E. Tijhuis, Floris Foijer, Andreas Villunger

## Abstract

Interference with microtubule dynamics in mitosis activates the spindle assembly checkpoint (SAC) to prevent chromosome segregation errors. The SAC induces mitotic arrest by inhibiting the anaphase-promoting complex (APC) via the mitotic checkpoint complex (MCC). The MCC component MAD2 neutralises the critical APC cofactor, CDC20, preventing mitotic exit. In cancer cell lines, this can provoke apoptosis involving pro-apoptotic BCL2 family members BIM and NOXA while BCL2 overexpression blocks mitotic cell death, facilitating SAC adaptation. However, the consequences of apoptosis after SAC perturbation in vivo are unclear. By conditional MAD2 overexpression across tissues in mice, we observed that chronic SAC activation triggers bone marrow aplasia and intestinal atrophy. While myelosuppression was tolerated, gastrointestinal atrophy was detrimental. Remarkably, co-deletion of Bim/Bcl2l11, but not Bid, Puma/Bbc3 or Noxa/Pmaip, prevented developing gastrointestinal syndrome caused by chronic SAC activity, identifying BIM as rate-limiting for mitotic cell death in the gastrointestinal epithelium. In contrast, only BCL2 overexpression but none of the BH3-only protein deficiencies tested could mitigate myelosuppression, highlighting tissue and cell type-specific survival dependencies in response to SAC perturbation in vivo.

## INTRODUCTION

During mitosis, genetic information needs to be propagated evenly between daughter cells. Mistakes in chromosome segregation and subsequent aneuploidy can harm cell fitness and organismal health^1, 2^. Several checkpoints monitor this process to regulate the fidelity of cell division. During mitosis, the spindle assembly checkpoint (SAC) is critical for adequately segregating sister chromatids into daughter cells^3, 4^. The main executioner of the SAC, the mitotic checkpoint complex (MCC), consists of BUBR1, BUB3, CDC20 and MAD2^5, 6^. During spindle formation in early mitosis, MAD2, encoded by *MAD2L1*, is conformationally activated by kinetochore-bound MAD1-MAD2, which converts inactive soluble open MAD2 conformation into its active closed form able to bind and neutralise CDC20. The MCC then binds to and prevents the full activation of the anaphase-promoting complex/cyclosome (APC/C), which is required for the ubiquitination-initiated degradation of securing and cyclin B1, which are required for sister chromatid separation and exit from mitosis, respectively. The SAC becomes inactivated upon bipolar chromosome attachment, which causes the depletion of MAD1-MAD2 from properly microtubule-attached kinetochores. Thus, the SAC monitors spindle formation and keeps cells in prometaphase until all chromosomes are correctly attached to the mitotic spindle^5, 7–9^.

Cell culture experiments suggest that mitotic cell death can be triggered if mitosis is prolonged by preventing SAC satisfaction in due time^10^. Alternatively, cells may escape prolonged mitosis by a process called “mitotic slippage” or “checkpoint adaptation”, triggered by the gradual non-canonical degradation of Cyclin B by the APC/C, allowing cell survival^11–14^. Slippage is frequently associated with increased ploidy and deregulated centrosome numbers, causing a substantial fraction of these cells to undergo senescence or cell death^15–17^. However, some poly- or aneuploidy cells can thrive and re-enter the cell cycle. This is facilitated by centrosome clustering, particularly when the function of the *p53* tumour suppressor gene is impaired or lost^18, 19^. Such features enable chromosomal instability (CIN) and aneuploidy and are frequently found in cancer^20^.

Complete loss of MAD2, or other MCC proteins leading to SAC deficiency, are embryonic lethal in mice, while hypomorphic alleles can cause microcephaly, premature ageing phenotypes and cancer predisposition syndromes^21–23^. In humans, to name a few examples, mutations of the SAC components cause mosaic-variegated aneuploidies ^24^, with germline mutations in *BUB1* linked to microcephaly^25^ and mutations in *MAD1L1* causing multiple malignancies^26^. Together, these findings underpin the importance of the mitotic spindle assembly checkpoint in development and tumour suppression. Of note, even partial loss of MAD2 function leads to premature degradation of Securin and separation of the sister chromatids, causing aneuploidy and polyploidy in cell lines and animal models. Consistently, haploinsufficiency accelerates T cell lymphomagenesis in a *p53*-deficient background and lung cancer progression in preclinical cancer models^27^.

Consistent with the mode of MAD2 action, overexpression can hyperactivate the SAC and lead to polyploidy, lagging chromosomes, and aneuploidy^28–30^. Consequently, tumourigenesis was accelerated by MAD2 overexpression together with MYC in blood cells or mutated KRAS in the lung epithelium^28, 29^. However, upon MAD2 overexpression, increased cell death rates were also noted, a phenomenon recently linked to FOXM1 transcription factor expression levels^31^. This may account for delayed tumour onset reported upon MAD2 overexpression in a *Her2-*driven mouse model of breast cancer^32^. While SAC genes, including *Mad2L1,* are rarely found mutated in human cancers, overexpression is observed more frequently (summarised by Simonetti, Bruno, Padella, Tenti and Martinelli ^33^). Besides ovarian cancer, increased MAD2 protein expression correlates with increased mortality and cancer recurrence in humans ^34^.

A series of in vitro studies using model cell lines have shown that the BCL2 protein family is critically involved in regulating mitotic cell death and cell death after slippage^11, 14^. The BCL2 family controls mitochondrial apoptosis. It comprises a set of anti-apoptotic molecules, including BCL2 itself, BCLX or MCL1, as well as BH3-only proteins that act as pro-apoptotic stress sensors (e.g. BIM, BID and NOXA), along with apoptotic effectors, BAX and BAK.

Later are required for pore-formation at the outer mitochondrial membrane, kick-starting a pro-apoptotic signalling cascade^35, 36^. During severe mitotic delays, the pro-survival BCL2-family member MCL1 acts as a molecular timer, where its NOXA-dependent degradation facilitates BIM-induced cell death in cancer cell lines^37^. In addition, the BH3-only proteins BID^38^ and BMF^31^ have been implicated in mitotic cell death and may act complementary to BIM and/or NOXA in contexts or cell types that still need to be defined. Loss of MCL1 expression renders epithelial cancer cells highly dependent on anti-apoptotic BCLX. This represents a therapeutic vulnerability that can be targeted by so-called “BH3 mimetics” that exploit the mode of action of BH3-only proteins, leading to BAX/BAK activation^39, 40^. Notably, mRNA expression of the BH3-only protein *NOXA* can be an independent survival predictor in human breast cancer patients treated with microtubule-targeting agents^41^.

While compelling, whether the same molecular players mediating cell death in response to spindle poisons in cancer cell lines ex vivo are also critically involved in the cellular response to chronic SAC deregulation in vivo was, to the best of our knowledge, never tested. Hence, we exploited a mouse model allowing conditional overexpression of the MCC component MAD2 across tissues^29^. Using this system, we aimed to address whether mitochondrial apoptosis limits the survival of cells experiencing chronic SAC activity to prevent aneuploidy and CIN-related pathology.

## RESULTS

### MAD2 overexpression triggers mitotic delay and cell death in haematopoietic cells

First, we assessed the impact of aberrant MAD2 expression on cell survival of haematopoietic cells. Herein, we used mouse bone marrow, immortalised with the homeobox gene *HoxB8*^42, 43^, to create SCF-dependent myeloid progenitors of neutrophils (dubbed PN) or multi-potent progenitor cells that depend on FLT3 ligand (dubbed PF), respectively. HoxB8-PN or HoxB8-PF cells were derived from mice, allowing overexpression of MAD2 upon Doxycycline (Dox) addition. In this model, N-terminally HA-tagged murine *Mad2L1* was knocked into the *Col1a* locus and controlled by the reverse tetracycline transactivator (rtTA) expressed from the *Rosa26* locus. This allows near ubiquitous transgene expression in response to Dox-treatment^32^. Additionally, we crossed-in a human BCL2 transgene, driven by the *Vav-*gene promotor, active in all haematopoietic cells^44^. Transgenic animals and bone-marrow-derived cell lines are referred to as: R (*R26rtTA*), MR (*Mad2/R26rtTA*), or MR2 (*Mad2/R26rtTA/BCL2*) throughout the text and figures. In some settings, these mice or cell lines also carried a Dox-responsive GFP-reporter, targeted to the second *Col1a* allele^45^. The *HA-Mad2* allele was maintained in a hemizygous state. Hence, as an example, primary MRG2-derived bone marrow cells harbour four transgenes (*Mad2/R26rtTA/GFP/BCL2*) that were eventually immortalised using a HoxB8-encoding retrovirus^46^.

Then, we induced MAD2 expression by Dox treatment in vitro and monitored transgene levels over time using an anti-HA antibody. Both bone marrow-derived cell lines, MR and MR2, showed increasing MAD2 levels in a dose- and time-dependent manner (Figure 1A; S1A). Constitutive overexpression of transgenic *BCL2* was confirmed in MR2 cells using a human BCL2-specific antibody, while endogenous BCL2, BCLX and MCL1 levels were comparable between both genotypes (Figure 1A; S1A, B). As noted before, high human BCL2 levels also allowed cells to tolerate higher levels of BIM (Figure 1A;^47, 48^). Interestingly, BIM levels dropped in response to Dox-treatment-induced MAD2 expression, best visible in BCL2 transgenic cells after 8 and 12h (Figure 1A). This likely reflects its increased turnover by the APC in cells experiencing mitotic delay or arrest^49, 50^.

**Figure 1:**
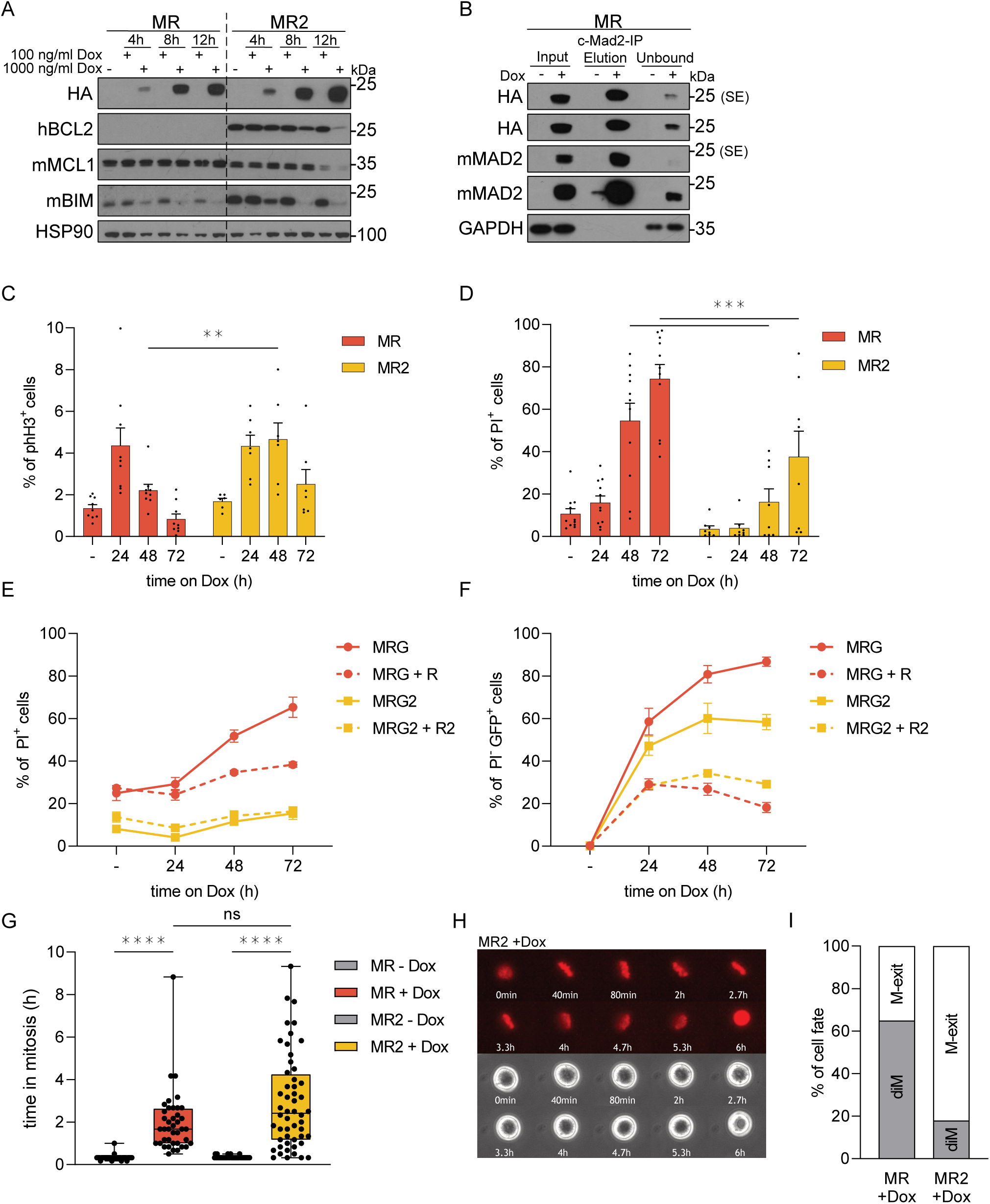
MAD2 overexpression triggers mitotic delays and BCL2-regulated cell death. HoxB8-PN cells (PNs) were generated from bone marrow of mice of the indicated genotypes and treated with Doxycycline for the indicated time points or 24h, if not stated otherwise. **(A)** PNs were harvested for immunoblot analysis with indicated antibodies after Mad2-transgene induction with 100/1000ng/ml Doxycycline. **(B)** Input, elution and unbound fractions from immunoprecipitation analysis to detect MAD2 in its closed (active) conformation. **(C)** PNs were fixed with ethanol and analysed by flow cytometry to quantify cells in mitosis staining positive for phH3^+^; MR (n=4-5), MR2 (n=3-4), two independent experiments. **(D)** HoxB8-PF cells were stained with PI and viability was analysed by flow cytometry: MR (n=4-5), MR2 (n=3-4), two independent experiments. **(E, F)** PNs were cultured alone or mixed 50:50 with transgene-negative cells in the presence or absence of Doxycycline and analysed for GFP expression and viability by flow cytometry; MRG (n=6), MRG+R (n=11), MRG2 (n=2) MRG2+R2 (n=6). (**G**) PFs were subjected to live-cell imaging to evaluate time spent in mitosis, from chromosome condensation until change of cell fate, with or without MAD2 induction. Min to max, Kruskal-Wallis-test; MR-Dox (n=40), MR+Dox (n=40), MR2-Dox (n=40), MR2+Dox (n=52). (**H**) Representative live-cell imaging time series of an MR2 transgenic cell after MAD2 induction, eventually dying in mitosis (DNA stained with SPY650-DNA & phase contrast). **(I)** Cell fate of cells shown in G, diM= death in mitosis, M-exit=cells exiting mitosis.

Next, we conducted immunoprecipitation experiments to demonstrate that the increase in MAD2 ends in increased levels of active closed MAD2^7, 9^. Using a conformation-specific MAD2 antibody, we confirmed excess active MAD2 upon Dox addition (Figure 1B). Increased levels of closed MAD2 showed a strong correlation with a higher percentage of phospho-histone H3 positive (phH3+) HoxB8-PN cells. This correlation was monitored using flow cytometry analysis and is indicative of mitotic delays or arrests in the cells (Figure 1C; S1C). Simultaneous viability analysis showed an increase in the percentage of MR cells being propidium iodide positive (PI^+^), indicating increased levels of cell death in cultures upon Dox-induced MAD2 overexpression (Figure 1D; S1D). The percentage of PI^+^ cells was strongly reduced in the presence of a BCL2 transgene (Figure 1D; S1D). Incucyte-based time-lapse microscopy analysis of MR and MR2 cells cultured in the presence of PI underlined the beneficial effect of BCL2 overexpression (Figure S1E). However, this effect vanished over time, indicating secondary necrosis in culture (Figure 1D).

We also exploited the Dox-responsive GFP-reporter as a surrogate marker of MAD2 expression in living cells by flow cytometry. GFP expression was readily detectable by flow cytometry analysis and on the protein level by immunoblotting (Figure S1F, G). We exploited this system to perform a competition assay comparing MAD2 transgenic cells that turn green upon Dox addition (MRG) and rtTA single transgene-positive control cells (R), expressing neither the GFP-reporter nor MAD2. In MRG cells, increased GFP expression correlated with increased cell death, as measured by PI uptake; BCL2 overexpression again blocked cell death in MRG2 cells (Figure 1E). When MRG cells were mixed in a 1:1 ratio with control cells (R), we observed that the latter rapidly outcompeted the MAD2 overexpressing cells, as inferred by the rapid plateau in viable GFP^+^ cells (Figure 1F). Notably, BCL2 transgenic cells failed to perform substantially better in this competition assay, indicating they may no longer proliferate at comparable rates (Figure 1F).

We performed live cell imaging to better understand cell fate in the presence of BCL2. We could observe that MAD2 overexpressing cells spend more time in mitosis than non-induced cells, ending primarily in mitotic cell death without cytokinesis (Figure 1G, H). As expected, cells arrested largely at metaphase, indicating SAC hyperactivation despite successful spindle formation. The duration of the mitotic arrest was not altered once BCL2 was overexpressed. Importantly, however, we noticed that the presence of BCL2 reduced death in mitosis (diM) from about 60% to 20% in MR2 cells, paralleled by an increase in cells managing mitotic (M)-exit (Figure 1I; movies S1-4).

As we noted reduced cell death and increased mitotic exit rates in BCL2 transgenic cells, we hypothesised that such cells may display signs of CIN. Therefore, we performed single-cell whole genome sequencing (scWGS) at different time points after induction of MAD2 overexpression. Flow cytometry was used to confirm the effects of MAD2 overexpression on cell cycle progression via phH3-staining and cell survival using sub-G1 analysis before sequencing analyses (Figure S2A, B). Surprisingly, we neither detected karyotypic abnormalities in the absence or presence of Dox in R, R2, MR or MR2 cells nor within MR2 cells still alive after 3 or 7 days (Figure S2C, D). This indicates that MR cells die in mitosis; those that survive due to exogenous BCL2 overexpression eventually must manage to inactivate SAC signalling, as we failed to observe aneuploidy or polyploidy despite MAD2 overexpression.

In summary, overexpression of MAD2 leads to mitotic delay and cell death in *HoxB8*-immortalized haematopoietic progenitor cells. However, MAD2 overexpression does not trigger viable karyotypic changes in tissue culture. The pro-survival protein BCL2 can reduce cell death induced by MAD2 overexpression. This may give cells enough time to terminate MAD2-induced SAC signalling and complete mitosis, as no karyotypic abnormalities were noted in scWGS analyses.

### Systemic overexpression of MAD2 can cause premature lethality

Next, we aimed to interrogate the consequences of MAD2 overexpression on the haematopoietic system in vivo by putting R and MR mice on Dox-containing diet. Unexpectedly, we noted that total-body transgenic mice rapidly lost body weight and 4/6 animals became clearly moribund within 5 days of Dox treatment (Figure 2A). Bone marrow cellularity, however, was not changed within this period (Figure 2B). A time-dependent increase of phH3^+^ cells within the bone marrow was noted from day 1 onwards, documenting SAC activation and indicating mitotic delays upon MAD2 overexpression (Figure 2C). Along this line, we detected increased cell death of bone marrow cells in situ from day 3 onwards (Figure 2D). Curiously, the percentage of LSK cells increased significantly upon MAD2 overexpression (Figure 2E, F). We also noted an accumulation of common lymphoid and common myeloid progenitors (CLP/CMP) upon SAC perturbation on day 5 (Figure S3A, B). The accumulation of LSK, CLP and CMP cell fractions in the bone marrow suggested delayed maturation or increased mobilisation due to the loss of more mature blood cell types. This aligned well with the significantly reduced rapidly dividing lymphocyte progenitors in the bone marrow and thymus. While the percentage of total B cells was not perturbed in the bone marrow (Figure 2G), we noted a clear decrease in cycling early pro/pre B cells, leading to a relative increase of mature recirculating B cells on day 5 (Figure 2H, I). Along similar lines, we observed a substantial reduction of thymic cellularity and atrophy upon MAD2 overexpression (Figure 2J). Immature double positive (DP) thymocytes were most affected, leading to a relative increase in the percentage of CD4^+^ and CD8^+^ single positive and double negative (DN) cells (Figure 2K, L). Granulocytes showed a mild increase within the myeloid compartment in the bone marrow, whereas Mac1^+^Gr1^-^ monocytes/macrophages decreased over time (Figure S3C, D). Consistent with their resting state, mature T and B lymphocytes in the spleen (Figure S3E-I) and lymph nodes were unchanged (Figure S3J-L). Erythroblasts mildly decreased in the bone marrow but stayed unaffected in the spleen (Figure S3M-O).

**Figure 2:**
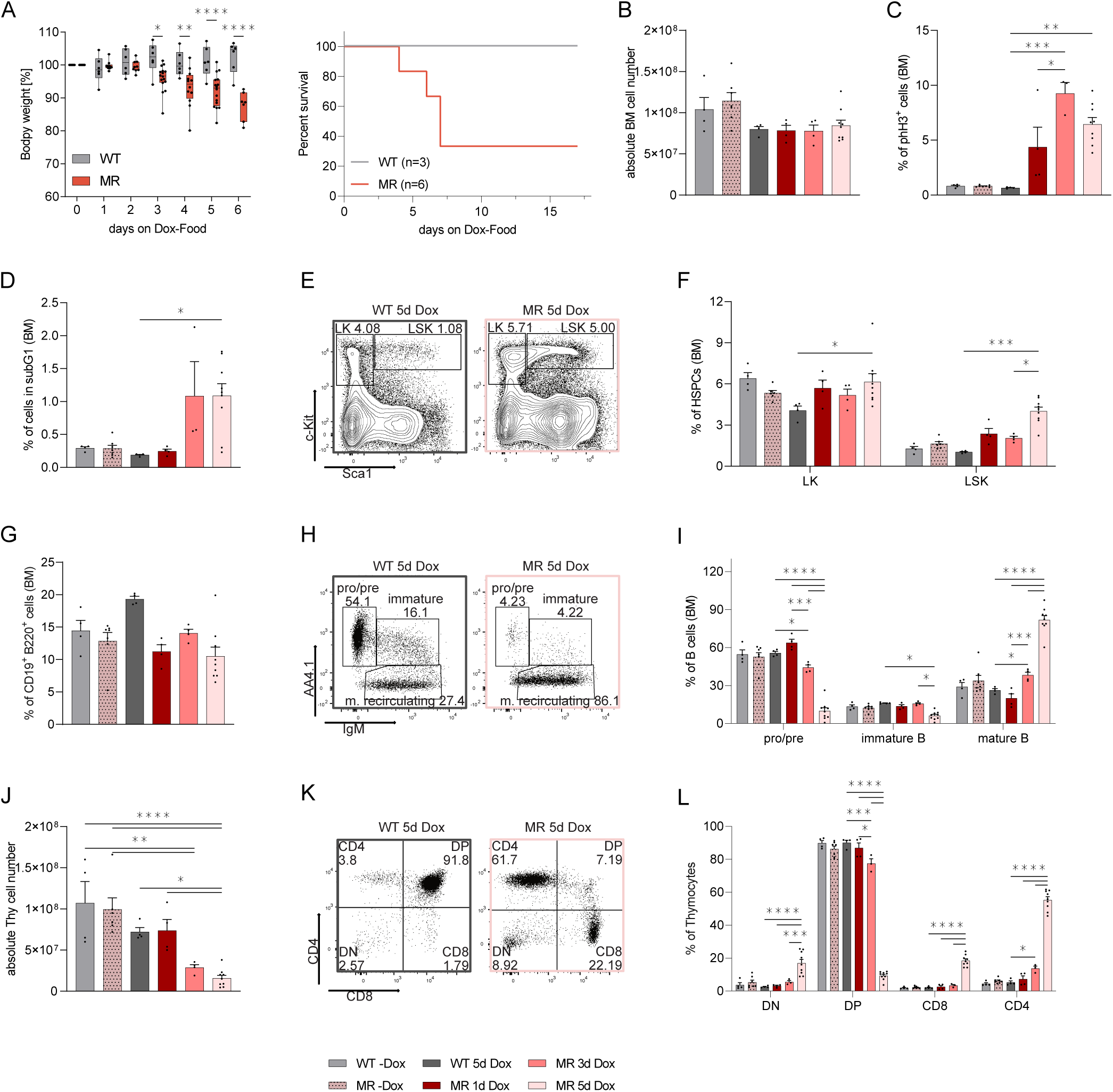
Systemic MAD2 overexpression causes rapid myelosuppression and wasting syndrome. **(A)** Body weight and Kaplan-Meier analysis of WT (R) and Mad2 transgenic animals (MR) treated with Doxycycline, R (n=3) MR (n=6). **(B)** Bone marrow (BM) cellularity of R and MR mice (both tibiae and femora) on Dox-containing food for the indicated time points. **(C)** Flow cytometric analysis of mitotic phH3^+^ bone marrow cells, **(D)** quantification of apoptotic BM cells by subG1 analysis. **(E)** Gating strategy used in flow cytometric analyses of HSPCs quantified in (F). **(F)** Percentage of LK (Lin^-^ckit^+^Sca1^-^) and LSK (Lin^-^ckit^+^Sca1^-^) in BM. **(G)** BM suspensions stained for total B (CD19^+^B220^+^) cells. **(H)** Gating strategy used in flow cytometric analyses of B cell subsets quantified in (I). **(I)** BM stained for pro/pre (AA4.1^+^IgM^-^), immature (AA4.1^+^IgM^+^) and mature recirculating (AA4.1^-^IgM^+^) B cells. **(J)** Thymic (Thy) cellularity of R and MR mice on Dox food for indicated time points. **(K)** Gating strategy used in flow cytometric analyses of thymocytes quantified in (L). **(L)** Thymocyte subsets assessed by CD4 and CD8 cell surface marker staining: DN (CD4^-^CD8^-^), DP (CD4^+^CD8^+^), single positive (CD4^+^CD8^-^) or (CD4^-^CD8^+^) cells. WT-Dox (n=4), MR-Dox (n=7), WT 5d Dox (n=4), MR 1d Dox (n=4), MR 3d Dox (n=4), MR 5d Dox (n=9).

Together, our data suggests that chronic SAC activation interferes with haematopoiesis by causing the loss of highly proliferative T and B cell progenitors. This leads to the mobilisation of HSCPs in the bone marrow and a subsequent increase in CMP and CLPs attempting to replenish haematopoiesis.

### MAD2 overexpressing haematopoietic stem and progenitor cells show reduced reconstitution potential

To study the impact of MAD2 overexpression on the ability of the haematopoietic stem and progenitor cell pool (HSPCs) to salvage the loss of developing leukocytes, we isolated the bone marrow from MAD2 transgenic animals (MR). We tested its colony-forming potential in methylcellulose in the presence or absence of Dox. IL-7 was added to promote pre B cell colony forming units (CFU-B), M-CSF for macrophage (CFU-M) and G-CSF for granulocyte (CFU-G) colony formation, respectively. We observed that pre B and myeloid colony formation was drastically impaired upon Dox-treatment. At the same time, untreated *Mad2* transgenic bone marrow could form colonies at frequencies similar to wild bone marrow (Figure 3A). Thus, MAD2 overexpression limits the clonogenic potential of freshly isolated haematopoietic stem and progenitor cells in vitro.

**Figure 3:**
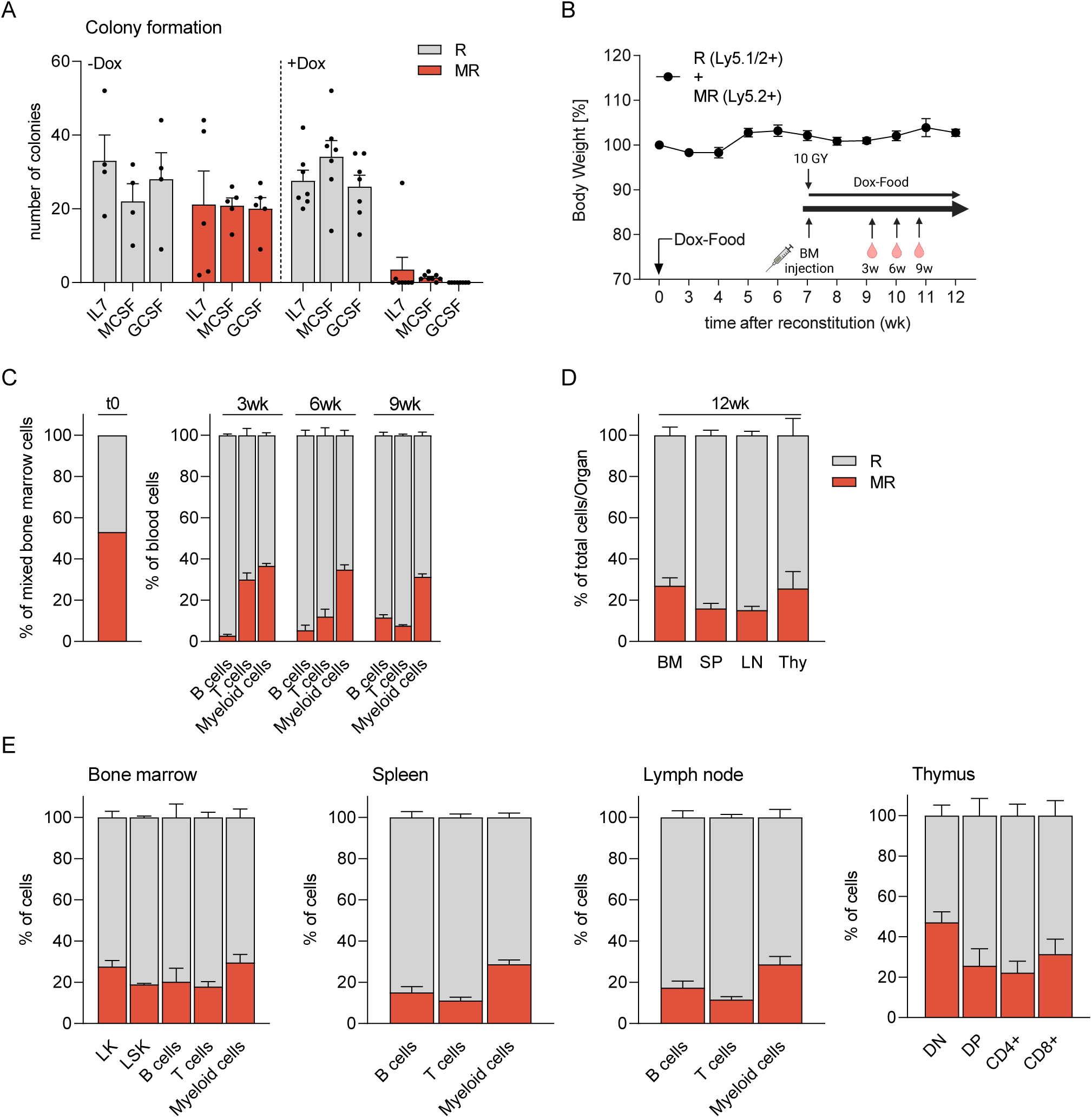
MAD2 overexpression perturbs fitness of haematopoietic stem and progenitor cells (HSPCs). **(A)** Colony formation potential of MAD2 (MR) and rtTA (R) transgenic bone marrow cells in methylcellulose assays in the presence (+) or absence (-) of Doxycycline; R+Dox (n=7), MR+Dox (n=8), R-Dox (n=4), MR-Dox (n=5). **(B)** Scheme of bone marrow reconstitution experiments and body weight analysis of chimaeric animals receiving a 50:50 mix of R (Ly5.1/2^+^) and MR (Ly5.2) BM cells. **(C)** The degree of chimaerism of peripheral blood cells was assessed by flow cytometry over time and compared to time zero (t0). Peripheral blood was sampled on indicated time points after reconstitution and diet change to analyse the content of B (CD19^+^B220^+^), T (CD4^+^&CD8^+^), and myeloid cells (CD11b^+^); percentages of R (Ly5.1/2^+^) and MR (Ly5.2^+^) cells within each cell type are shown. **(D)** The degree of chimaerism in haematopoietic organs of reconstituted animals was analysed after 12 weeks by flow cytometric analysis (BM: Bone marrow; SP: spleen; LN: lymph node; Thy: Thymus). **(E)** BM, SP, LN and Thy were analysed after 12 weeks for the % of R and MR cells in different haematopoietic cell types: LK (Lin^-^Sca1^-^cKit^+^) and LSK (Lin^-^ Sca1^+^cKit^+^), B (CD19^+^B220^+^), T (CD4^+^ and CD8^+^), myeloid cells (CD11b^+^), DN (CD4^-^CD8^-^), DP (CD4^+^CD8^+^), T helper (CD4^+^) and T cytotoxic (CD8^+^) cells; (n=4).

We next investigated the consequences of aberrant MAD2 expression on the haematopoietic system in vivo in a competitive reconstitution setting to avoid potentially detrimental bone marrow failure. Thus, we isolated isogenic *Mad2* transgenic (MR, Ly5.2^+^) and *rtTA* (R, Ly5.1/2^+^) bone marrow (Figure S4A) and injected a 50:50 mix into lethally irradiated recipient (Ly5.1^+^) mice. Bone marrow chimaeras were fed Dox food during haematopoietic reconstitution. The animals showed stable body weight throughout the 12-week observation period (Figure 3B). Analysis of peripheral blood showed rapid displacement of the *Mad2* transgenic (MR) Ly5.2^+^ cells by the *rtTA* single-transgenic (R) Ly5.1/2^+^ leukocytes. B cells appeared to be most affected, followed by T cells that decreased over time, while myeloid cells were more resilient (Figure 3C). These results suggest varying susceptibility of different leukocyte subtypes or their respective progenitors to SAC perturbation. In line with our peripheral blood cell analysis, bone marrow, spleen, lymph nodes, and thymus contained mostly haematopoietic cells originating from *rtTA* bone marrow (R) when analysed 12 weeks after reconstitution (Figure 3D; S4B). Flow cytometry analysis documented reduced percentages of Lin^-^Sca1^+^cKit^+^ (LSK) bone marrow cells, a population of cells enriched in HSCPs, and Lin^-^cKit^+^ (LK) bone marrow cells, containing committed haematopoietic progenitors. MR-derived myeloid cells and mature T and B cells were also outnumbered (Figure 3E; S4C, D). Together, these findings document reduced fitness of MAD2 overexpressing haematopoietic stem and progenitor cells in vivo, likely leading to a reduced output of leukocytes. However, the impact of chronic SAC activation on leukocyte homeostasis remained uncertain.

### MAD2 overexpression impairs leukocyte homeostasis but fails to induce malignancies

Next, we investigated the consequences of SAC perturbation on haematopoiesis in the presence or absence of a potent cell death inhibitor, i.e. transgenic BCL2. We generated pure bone marrow chimaeras using RG, MRG, RG2 and MRG2 animals as bone marrow donors (Figure 4A). After reconstitution for 12 weeks, animals were set on a Dox-containing diet to induce *Mad2* transgene expression that was monitored using GFP induction as a surrogate marker. Similar to the 50:50 reconstitution setting, the expression of Ly5.2^+^ cells allowed us to discriminate between donor (Ly5.2^+^) and recipient (Ly5.1^+^) cells. Peripheral blood analysis before and after Dox food confirmed the successful reconstitution of mice with >90% of Ly5.2^+^ donor cells of various genotypes. This phenotype remained consistent over time with comparable percentages of white (WBC) and red blood cell counts (RBC) in RG and MRG recipients, as well as normal haematocrit (HCT) values. As expected, the presence of the *BCL2* transgene initially increased white blood cell counts^44, 51^ in RG2 and MRG2 recipients. Still, RBC and HCT were comparable to that seen in RG and MRG reconstituted animals (Figure S5A, B). The distribution of B and T lymphocytes and myeloid cells in the peripheral blood did not change significantly over Dox treatment (Figure S5C).

**Figure 4:**
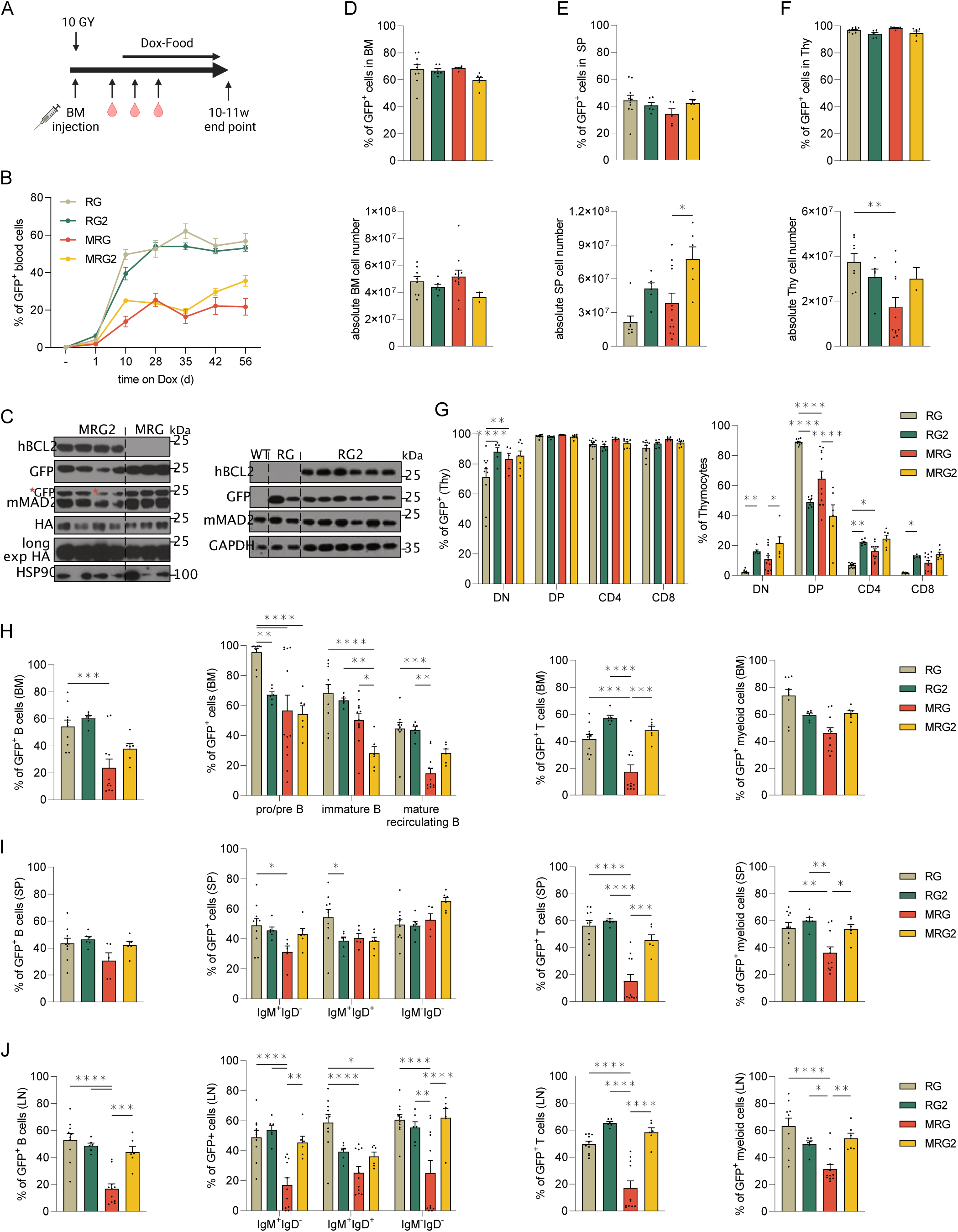
MAD2-induced myelosuppression is ameliorated by transgenic BCL2. **(A)** Scheme of bone marrow reconstitution analysis. **(B)** Total GFP expression in the peripheral blood of chimaeras of indicated genotypes was monitored over time. **(C)** Bone marrow of reconstituted animals was subjected to Western analysis using the indicated antibodies (*: persisting GFP signal after re-probing, WT: non-reconstituted mouse). Flow cytometric analysis of GFP expression and quantification of absolute cell counts in **(D)** bone marrow (BM, both femora and tibiae), **(E)** spleen (SP) and **(F)** thymus (Thy) of mice of the indicated genotypes 12 weeks after reconstitution and additional 10 weeks of Dox treatment. **(G)** Analysis of GFP expression of thymocyte subsets quantified in H. **(H)** Thymocytes were stained using CD4 and CD8-specific antibodies to identify CD4^-^CD8^-^ double-negative (DN), CD4^+^CD8^+^ double-positive (DP), CD4^+^CD8^-^ single positive helper and CD4^-^CD8^+^ cytotoxic T cell progenitors. GFP^+^ cells within **(H)** BM **(I)** SP and **(J)** LN (lymph node) were analysed for the presence of total B cells (CD19^+^B220^+^) and different B cell subsets: pro/pre (AA4.1^+^IgM^-^), immature (AA4.1^+^IgM^+^) and mature recirculating (AA4.1^-^IgM^+^), or IgM^+^IgD^-^, IgM^+^IgD^+^, IgM^-^ IgD^-^ B cells of total B cells; T (CD4^+^ and CD8^+^) and myeloid cells (CD11b^+^). RG (n=8-10), RG2 (n=6), MRG (n=5-11), MRG2 (n=2-6).

Notably, the fraction of GFP^+^ cells in the peripheral blood increased on day 1 after providing Dox food. About 60% of GFP^+^ cells were observed after 28 days in RG and RG2 recipients (Figure 4B), while animals reconstituted with *Mad2* transgenic bone marrow (MRG) showed a significantly lower percentage of GFP^+^ blood cells than controls (RG). Curiously, while MRG2 recipients showed a faster increase in the percentage of GFP^+^ cells, as seen on day 10, potentially due to impaired cell death, both MRG and MRG2 recipients reached a plateau of about 20% and 30% GFP^+^ cells, respectively (Figure 4B). This again indicated reduced fitness or proliferative capacity of MAD2 overexpressing blood cells aiming to replace resident leukocytes over time with BCL2 overexpression, providing a relatively modest advantage.

After 10 weeks of Dox treatment, mice were sacrificed for organ analysis. BCL2 and GFP-transgene expression were confirmed along with HA-MAD2 in the bone marrow of recipient mice by immunoblot (Figure 4C). The relative abundance of T, B and myeloid cells was broadly comparable across the different genotypes and organs, showing similar percentages of these populations in flow cytometric analyses. A notable exception was an increase in myeloid cell infiltrates into lymph nodes in response to MAD2 overexpression, suggesting sterile inflammation (Figure S5D-F). Most importantly, we observed that MAD2 expression from the *Col1a* locus was highly varied in different haematopoietic organs. The highest percentages of GFP^+^ cells were found in the thymus (Thy), followed by the bone marrow (BM) and the spleen (SP) (Figure 4D-F).

On the one hand, organ cellularity was similar in bone marrow across genotypes (Figure 4D). At the same time, BCL2 overexpression led to the expected increase in splenocytes^44, 51^ in the absence or presence of MAD2 (Figure 4E). On the other hand, MAD2 overexpression led to a selective reduction in thymic cellularity (Figure 4F), caused by a significant drop in CD4^+^CD8^+^ immature double-positive cells that was not seen on a BCL2 transgenic background (Figure 4G). BCL2 overexpression also led to a general redistribution of different thymocyte subsets ^44^, but this phenomenon was not affected by excess MAD2 (Figure 4G). Evaluating the leukocyte subset distribution within the GFP^+^ compartment, we noted that MAD2 overexpression led to a clear reduction of reporter-positive leukocyte cells. This reduction within bone marrow spleen and lymph nodes (LN) comparing MRG with RG recipients was buffered by BCL2 overexpression (MRG2 vs. RG2) (Figure 4H-J).

*Vav-BCL2* transgenic animals show severely reduced lifespan due to autoimmune disease or follicular lymphoma development, exacerbated in a bone-marrow reconstitution setting ^44, 52^. Therefore, we could not study the long-term consequences of cell death inhibition after MAD2 overexpression in this model. Instead, to monitor the spontaneous transformation potential of MAD2 alone, we followed a small cohort of seven MRG mice for one year on Dox food. Analysis of peripheral blood for the percentage of total Ly5.2^+^ cells showed a stable reconstitution with MRG bone marrow (Figure S6A). HCT, RBC and WBC in these animals were stable over time, as was the distribution of leukocytes (Figure S6B, C). The fraction of GPF^+^ cells in the blood increased gradually, reaching more than 90% after 12 months (Figure S6D). Interestingly, after one month, the percentage of GFP^+^ myeloid cells reached a plateau of about 70%. Turnover of B and T cells required more time, reaching >80% after the endpoint of our blood cell analysis (Figure S6E). A final analysis of primary and secondary haematopoietic organs after 12 months confirmed an increased percentage of phH3^+^ bone marrow, suggesting the *Mad2* transgene was still expressed and active in haematopoietic cells (Figure S6F). Consistently, MAD2 overexpression was well-detectable on protein level in bone marrow, spleen and thymus (Figure S6G-I). Organ cellularity and the composition of haematopoietic cells in bone marrow, spleen, lymph nodes (Figure S6J) and thymus (Figure S6K) were comparable to that of MRG animals analysed after 2 months (Figure 4). None of the animals did develop signs of malignant disease.

In summary, we noted clear signs of myeloablation in MRG recipients after transgene induction that were ameliorated by BCL2 overexpression. This suggests a role for mitochondrial apoptosis in response to chronic SAC perturbation in white blood cells. Moreover, due to variegated transgene expression, MAD2 overexpression in the haematopoietic system is tolerated with minimal impact on organ cellularity. Still, MAD2 overexpression appears insufficient to promote spontaneous malignancies within 12 months, likely due to increased apoptotic priming.

### Chronic SAC perturbation causes severe colitis and gastrointestinal syndrome

Our results in reconstitution experiments suggested that myelosuppression is limited and *Mad2* transgene expression variegated in blood cells, hence not causal for the rapid wasting of MR animals. Based on the similarities of the phenotype seen with radiation disease, we interrogated the gastrointestinal (GI) tract. GI tract integrity and barrier function rely on the constant and rapid replacement of short-lived gastrointestinal epithelial cells, shed off at the tip of villi and replaced by newly differentiating cells originating from the crypt-based stem cell and transient amplifying pool^53, 54^.

Histopathological assessment of the intestine revealed a severe colitis/enteritis phenotype in *Mad2* transgenic animals (Figure 5A, B), likely explaining the animals’ weight loss and rapid wasting. MAD2 transgene expression was verified in the GI tract after Dox addition by IHC (Figure 5C) and Western blot of total intestinal lysates (Figure S7A). We also confirmed increased phH3^+^ cells in the colonic and small intestinal crypts of *Mad2* transgenic animals, indicating the expected mitotic arrest and SAC activation (Figure 5C, D). This correlated well with increased active cleaved Caspase 3^+^ cells in both the colon and the small intestine, documenting increased cell death (Figure 5C, E) and elevated p21 levels (Figure 5C; S7A). Evaluating expression levels of different BCL2 family proteins indicated lower levels of BCL2 and BIM, a reported APC substrate^49, 50^, in response to Dox treatment, while levels of BCLX and MCL1 appeared unchanged (Figure S7A).

**Figure 5:**
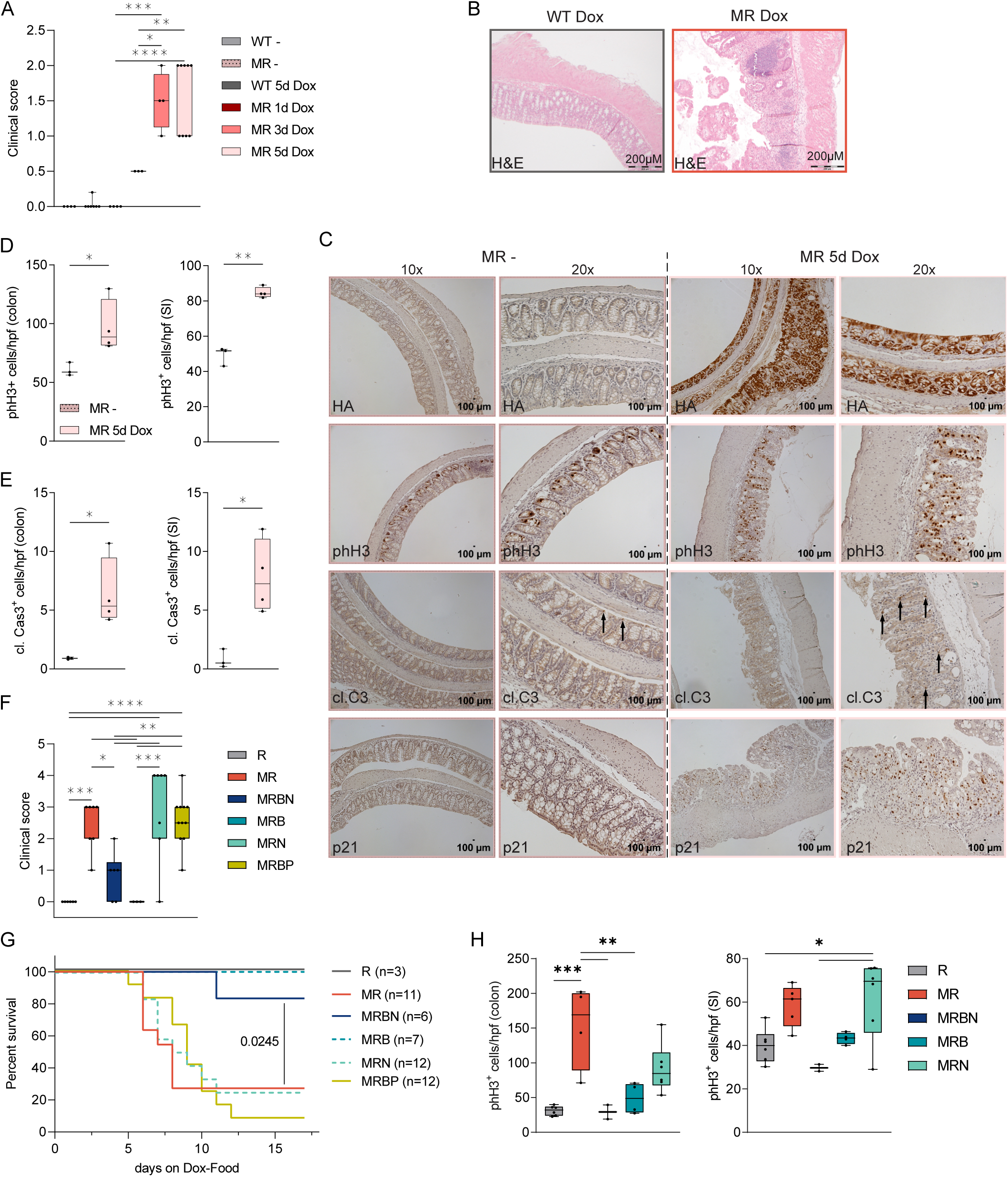
Systemic MAD2 overexpression causes detrimental atrophy in the GI-tract ameliorated by loss of BIM. **(A)** Clinical score of the intestine of animals on regular diet or Dox-containing food. WT-(n=3), MR-(n=7), WT 5d Dox (n=3), MR 1d Dox (n=3), MR 3d Dox (n=4), MR 5d Dox (n=9). **(B)** H&E staining of paraffin-embedded sections from swiss rolls generated from healthy and terminally sick animals. **(C)** Representative pictures of MR animals on regular diet (MR-) and 5d on Dox food (MR 5d Dox), stained for HA, phH3, C3 (active Caspase 3) and p21 (10x and 20x magnification are shown). Quantification in colon and small intestine (SI) stained for **(D)** mitotic cells (phH3^+^) and **(E)** cleaved C3^+^ cells. MR-(n=3), MR 5d Dox (n=4). **(F)** Clinical score of animals with the indicated genotypes at the time of sacrifice (>17% loss of body weight) or after 17 days. **(G)** Kaplan-Meier analysis of mice overexpressing Mad2 in the absence or presence of different BH3-only proteins. **(H)** Quantification of colon and small intestine (SI) stained for mitotic cells (phH3^+^).

We hypothesised that the deletion of pro-apoptotic BIM together with NOXA, both needed for mitotic cell death in epithelial cancer cells^37, 55^, might ameliorate the wasting syndrome noted in *Mad2* transgenic mice by allowing GI-tract stem cells experiencing mitotic delays to evade cell death. Indeed, we could observe that the loss of *Bim* and *Noxa* damped the colitis phenotype (Figure 5F) and extended the lifespan of *Mad2* transgenic (MRBN) mice (Figure 5 G; S7B). Wondering whether the deletion of both BIM and NOXA is necessary for this survival advantage, we also analysed MAD2 overexpressing animals lacking only *Noxa* (MRN) or *Bim* (MRB). Of note, MRN mice developed severe colitis followed by a wasting syndrome similar to MAD2 overexpressing MR mice, while MRB animals showed no phenotype (Figure 5F, G). Surprisingly, we observed a reduction in the number of phH3^+^ cells in the absence of *Bim* in MRBN and MRB mice (Figure 5H) despite strong HA expression in the intestine (Figure S7B). This suggests that these cells may escape mitotic arrest and death in mitosis. Importantly, the co-deletion of *Bid* and *Puma/Bbc3* (MRBP), two other pro-apoptotic BH3-only proteins, could not rescue the colitis/enteritis phenotype (Figure 5F) nor extend the lifespan upon aberrant MAD2 expression (Figure 5G). Taken together, our data suggest that delays in mitotic cell death caused by loss of BIM can overcome the detrimental effects of impaired SAC proficiency in the GI tract allowing escape from mitotic arrest.

### Loss of BH3-only proteins fails to prevent MAD2-induced myelosuppression

To evaluate the contribution of mitotic cell death to the observed myelosuppression, we also investigated the impact of BH3-only protein deletion on blood cell formation and survival. Animals were analysed once they became moribund or on day 17 when remaining unaffected by Dox treatment (Figure 5G).

Similar to our observations in the GI tract, loss *Bim* caused a decrease in the fraction of mitotic phH3^+^ bone marrow cells after Dox treatment in MRB and MRBN mice (Figure 6A). Bone marrow cellularity and the percentage of HSPCs were not significantly changed within the treatment window (Figure 6B, C). However, loss of *Bim* and *Noxa* did not translate into rescuing progenitor B cells in bone marrow (Figure 6C) or T cell progenitors in the thymus (Figure 6D; S7D).

**Figure 6:**
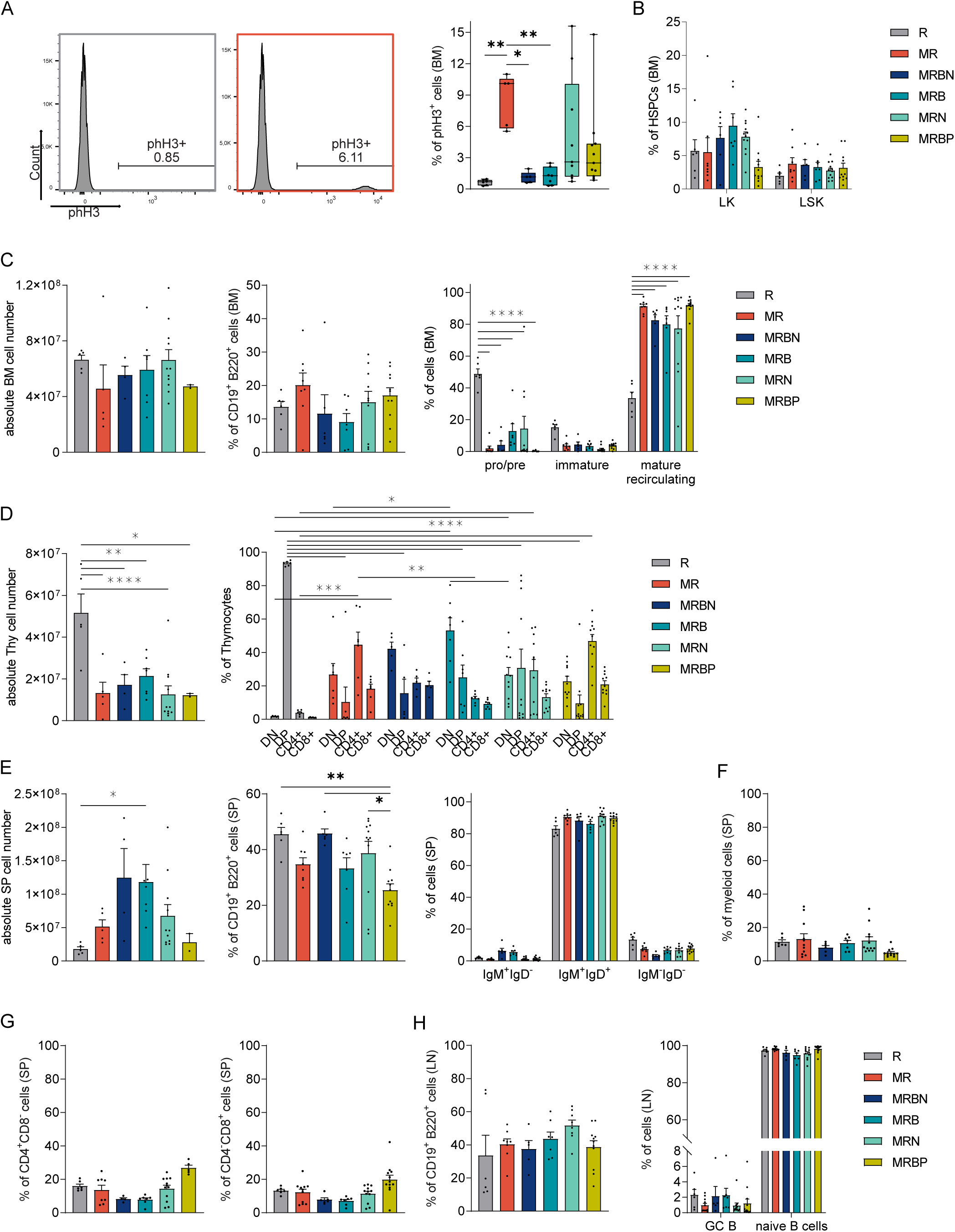
MAD2-induced myelosuppression is independent of BIM and NOXA. **(A)** Gating strategy used in flow cytometric analyses of BM cells, stained with an antibody specific for phH3 in mice of the indicated genotypes and MAD2 induction by the addition of Dox-containing food. Bone marrow (BM) of WT, MR, MRBN (Mad2/rtTA/Bim^-/-^/Noxa^-/-^), MRB (Mad2/rtTA/Bim^-/-^), MRN (Mad2/rtTA/Noxa^-/-^) and MRBP (Mad2/rtTA/Bid^-/-^/Puma^-/-^) mice when terminally sick or at the end time point of 17 days on Dox food stained for mitotic (phH3^+^) and **(B)** LK (Lin^-^ckit^+^Sca1^-^) and LSK (Lin^-^ckit^+^Sca1^-^) cells. **(C)** BM cellularity (both femora and tibiae) and staining for B (CD19^+^B220^+^) cells and subsets: pro/pre (AA4.1^+^IgM), immature (AA4.1^+^IgM^+^) and mature recirculating (AA4.1^-^IgM^+^) B cells. **(D)** Thymic (Thy) cellularity and thymocyte subsets identified by immune-staining: DN (CD4^-^CD8^-^), DP (CD4^+^CD8^+^), single positive T helper (CD4^+^CD8^-^) and T cytotoxic (CD4^-^CD8^+^) cells. **(E)** Spleen (SP) cellularity and staining for B (CD19^+^B220^+^) cells defining IgM^+^IgD^-^, IgM^+^IgD^+^, IgM^-^IgD^-^ subsets, **(F)** myeloid (CD11b^+^) cells and **(G)** T cells (CD4^+^CD8^-^, CD4^-^CD8^+^). **(H)** Lymph node (LN) stained for B (CD19^+^B220^+^) cells, germinal centre (GC, CD38^int^CD59^+^) and naïve (CD38^int^CD59^-^) B cells. R (n=6), MR (n=5-8), MRBN (n=4-6), MRB (n=7) and MRN (n=10), MRBP (n=2-11).

Splenic cellularity and weight were significantly increased in MRBN and MRB animals (Figure 6E, S7E). However, this can be explained by the splenomegaly caused by the loss of *Bim* ^47^. No changes were observed in the distribution of IgM^+^IgD^+^ mature B cells, CD4^+^ or CD8^+^ T cells, as well as myeloid cells in the spleen (Figure 6E-G) or lymph nodes (Figure 6H; S7F, G). These findings contrast observations made in BCL2 overexpressing HoxB8 cells (Figure 1) and mice reconstituted with BCL2 transgenic bone marrow, where some degree of protection from the consequences of MAD2 overexpression was noted (Figure 3). In line with the above, the colony formation potential of *Mad2*-transgenic HSPCs was not restored by loss of BIM and NOXA, as tested in methylcellulose assays (Figure S8A). Moreover, the loss of both BH3-only proteins caused only a very modest delay in the cell of HoxB8-immortalized progenitors after MAD2 overexpression ex vivo, contrasting findings made with BCL2 overexpression (Figure 1; S1E, S8B-D). This suggests that additional apoptosis effectors next to BIM are at play to remove haematopoietic cells experiencing substantial mitotic delays in vivo.

## DISCUSSION

Defects in the spindle assembly checkpoint are linked to developmental defects, premature ageing phenotypes and malignant disease while targeting this checkpoint has been exploited to treat cancer patients for several decades^15, 56, 57^. Our understanding of systemic and cellular responses contributing to pathology, cancer treatment efficacy and side effects is still incomplete. MAD2 overexpression delays mitosis, fosters chromosome miss-segregation and aneuploidy in fibroblasts and accelerates MYC-driven and spontaneous tumour initiation in transgenic mice^29^. Of note, Sotillo, Hernando, Diaz-Rodriguez, Teruya-Feldstein, Cordon-Cardo, Lowe and Benezra ^29^ showed that the tumour spectrum observed in MAD2 overexpressing mice phenocopied those found in human cancer with high MAD2 levels and poor prognosis, such as hepatocellular and lung carcinomas, as well as diffuse large B cell lymphomas (DLBCL). However, the need for additional genomic alterations to allow tumour formation might explain the rather long tumour latency reported in this model^29, 33^. Moreover, the anti-tumourigenic effects of MAD2 overexpression have also been noted in *Her2*-driven breast cancer^32^.

Consistent with the above, we observed increased cell death *in vitro* in myeloid progenitor cells overexpressing MAD2 (Figure 1; S1). In line with this finding, we failed to note an increase in poly- or aneuploidy in HoxB8 model cell lines. Moreover, even though we could delay cell death by BCL2 overexpression, we did not detect increased levels of aneuploidy by scWGS (Figure S2). How chronic MCC signalling is overcome in these cells remains to be investigated. Increased levels or activity of p31coment may be involved^58, 59^, but lack of commercially available mouse-specific antibodies precluded us form investigating this possibility. Of note, cell death was still seen at later time points, suggesting secondary necrosis or the activation of alternative cell death pathways during ex vivo culture of BCL2 transgenic cells (Figure 1). Consistently, we also observed that MAD2 overexpression causes the loss of cycling lymphocyte progenitors (Figure 2) and limits the clonogenic potential of primary haematopoietic stem and progenitor cells (HSCP) in vitro and in vivo, best seen in competitive reconstitution assays (Figure 3). If this happened solely by the induction of apoptosis, as seen in our ex vivo culture, or whether the induction of cell cycle arrest and senescence may contribute could not be distinguished using these readouts. The use of fully reconstituted animals expressing MAD2 together with a BCL2 transgene, however, supports a role of cell death induction in leukocytes also in vivo, as consequences of MAD2 overexpression were ameliorated by blocking mitochondrial apoptosis (Figure 4). In light of these observations, the lack of spontaneous blood cancer development in animals overexpressing MAD2 in the haematopoietic system may not come as a surprise (Figure 4; S6). However, this contrasts with findings originally made in *Mad2* transgenic animals that occasionally developed lymphomas during their first year of life and hence survived transgene expression long term ^29^. One may speculate about differences in genetic background and the use of a different transactivator for transgene expression, i.e. *CMV-rtTA* vs. *R26rtTA*, used here, that are known to differ in strength and cell type-specificity ^60^. Moreover, surviving cells with complex karyotypes might be cleared more effectively by NK cells in our model. The latter phenomenon was recently reported in cell line studies describing the killing of aneuploid cells by NK cells ex vivo^61, 62^.

Moreover, alterations in the non-haematopoietic compartment due to MAD2 transgene expression may add to the differences in tumour incidence. Consistent with this idea, acceleration and delays in tumour development were shown upon MAD2 overexpression. In a KRAS-driven mouse model of lung cancer, MAD2-induced CIN accelerated tumourigenesis and relapse after oncogene silencing^28^. Conversely, in a *Her2*-driven mouse model of breast cancer, MAD2 overexpression caused a delay in tumour onset but facilitated tumour persistence after oncogene withdrawal^32^. Cell type-specific differences in SAC impairment driving transformation in combination with different oncogenic drivers may likely explain these conflicting results.

Based on our reconstitution experiments using GFP as a reporter for MAD2 expression, we concluded that reduced fitness of MAD2 overexpressing HSCs and initial myeloid aplasia was overcome by transgene negative blood cells due to variegated transgene expression (Figure 4), allowing the long term survival of these mice. This phenomenon was apparently not pronounced in the GI tract stem cells, where transgene overexpression was detectable throughout all crypts in the small intestine and colon over extended periods with no signs of variegated expression patterns (Figure 5; S7). Consistently, gastrointestinal stem cells and the transient amplifying pool seemed to be the most affected cells by MAD2 overexpression and extended mitosis as an imbalance in MAD2 expression in either direction can cause prolonged SAC activation^27, 29^. Our finding is well in line with a recent report showing that timed conditional deletion of MAD2 in *Mad2l1^f/f^* mice carrying a *CRE^ERT2^* transgene triggers a similar phenotype with rapid weight loss, mitotic abnormalities and increased apoptosis in intestinal crypts, causing rapid animal death^63^.

Our study shows myelosuppression due to MAD2 overexpression is mainly limited to early cycling T and B lymphocyte progenitors. Accordingly, primary haematopoietic organs, bone marrow and the thymus were affected most strongly, with thymocytes and early pro/pre B cell progenitors affected most severely (Figure 2, 3). Moreover, we found an increase in the fraction of HSPCs in the bone marrow of *Mad2* transgenic mice (Figure 2), which indicates a mobilization response to compensate for the loss of T and B cell lymphoid progenitors. Consistently, we noted an accumulation of CMP and CLPs in the bone marrow (Figure S3). Alternatively, a block in differentiation may lead to an accumulation of HSPCs, fostering myelosuppression. Yet, based on all our findings, the former appears more compatible with the cell death-inducing effects of MAD2 overexpression in the HoxB8 model and the drop in early B and T cell progenitors.

In contrast, the number of mature T and B cells appeared normal in the reconstitution setting. Still, both were found to be reduced within the GFP^+^ fraction of cells (Figure 4), mimicking results from mixed bone marrow chimaeras (Figure 3). As those cells are already differentiated and do not proliferate unless challenged by antigens, the noted drop may reflect a loss of influx of new cells rather than deletion of mature lymphocytes. Yet, BCL2 overexpression ameliorated the loss of immature and mature lymphocytes within the reconstitution setting (Figure 4), suggesting that progenitors may be able to mature further in the presence of MAD2.

It is worth mentioning that mice that survived the systemic overexpression of MAD2 did not show such signs of myelosuppression at the time of analysis but were still diagnosed with colitis of lower grades. This suggests intestinal stem cells are more vulnerable to MAD2 overexpression and unable to escape transgene-induced effects unless cell death is perturbed. The most striking finding from our study is that we could increase the lifespan of *Mad2* transgenic animals by the co-deletion of the pro-apoptotic protein BIM and NOXA. Our follow-up experiments identified BIM as the most critical cell death effector, as the loss of several other BH3-only proteins tested, including the combination of BID and PUMA or NOXA alone, provided no noticeable survival advantage. This finding complements our previous in vitro studies, where we could show that upon extended mitotic arrest induced with microtubule targeting agents, anti-apoptotic MCL1 and NOXA are co-degraded, thereby releasing BIM that can initiate apoptosis ^37^. BIM is a potent pro-apoptotic protein, and it was shown that the deletion of BIM in mice causes lymphadenopathy but can also restore the detrimental effects of loss of BCL2, especially on mature B and T cell homeostasis^47^. As such, it was somewhat unexpected to learn that loss of BIM, alone or in combination with NOXA, failed to dampen myelosuppression and restore early lymphopoiesis in the thymus or bone marrow (Figure 6). Based on our observations using BCL2 overexpression in cultured cells (Figure 1; S8), we conclude that BIM may act in conjunction with other BH3-only proteins but not NOXA. Interaction with BMF^31^ or BID^38^, both implicated in mitotic cell death and noted before interacting with BIM in other pathological contexts^64, 65^, deserve to be tested. Alternatively, developing T and B cell progenitors may activate alternative cell death pathways when facing mitotic delays that do not rely on BIM or any of the combinations we tested. Consistently, cell type-dependent differences regarding the role of BIM in mitotic cell death have been reported^37, 59^. What stands out, however, is the observation that loss of BIM correlated with a reduction of phH3^+^ cells in the GI tract and bone marrow in situ (Figure 6A, S7C), contrasting findings in HoxB8 cells (Figure S8B). While indicating higher rates of MAD2 inactivation in both cases, this appears to enhance cell survival in the GI tract, but not the bone marrow. How the loss of BIM would affect adaptation is unclear. Still, similar to p31comet exerting undefined anti-apoptotic roles in mitotically arrested cells^59, 66^, BIM may restrain MCC function by yet-to-be-defined mechanisms.

In conclusion, our study identifies BIM as rate limiting for apoptosis induction in the gastrointestinal epithelium in response to SAC perturbation. In contrast, only BCL2 overexpression, rather than BH3-only protein deficiency, demonstrated the capacity to mitigate myelosuppression. These findings underscore the significance of tissue-specific survival dependencies in response to SAC perturbation. This aspect has potential implications for treatment strategies seeking to merge BH3 mimetics with spindle poisons or innovative inhibitors targeting the cell cycle machinery, such as MPS1, PLK1, or Aurora kinase inhibitors. The varying responses within different tissues may influence the therapeutic effectiveness and the potential side effects of such treatment regimens.

## Material and Methods

### Animals used and generation of bone marrow chimaeras

Animal experiments were performed in accordance with Austrian legislation (BMWF: 66-011/0106-WF/3b/2015 and GZ: 2021-0.222.1888). The generation of *Vav-BCL2* transgenic^44^, *Bcl2l11/Bim^-/-47^*, *Bbc3/Puma^-/-67^*, *Pmaip/Noxa^-/-67^*, *GFP*-reporter^45^ and *Mad2* transgenic animals^32^ were described before. All mice were maintained or backcrossed on a C57BL/6N genetic background or backcrossed for at least 6 generations in case of *Mad2* transgenic mice. Mice were housed in ventilated cages with nesting material and were maintained on a 12:12-h light:dark cycle.

Bone marrow reconstitutions were performed using C57BL6/N Ly5.1 recipient mice at the age of 8-10 weeks. Mice were irradiated with a single dose of 10Gy and reconstituted with 200µl PBS containing 4×10^6^ total bone marrow cells of Ly5.2 or Ly5.1/2 donor origin, injected via the tail vein. Mice were given Neomycin for 2-3 weeks in the drinking water before swapping to regular water. Mice were set directly (50:50 chimaeras) or 12 weeks after reconstitution on Dox food. Chimaeras were analysed after additional 10 weeks of transgene induction. Whole body MAD2 overexpressing mice were analysed at age 15-20 weeks indiscriminate of sex. Animals were examined once terminally sick (losing >17% of initial body weight) or after surviving for 17 days on Dox food. The genotype was blinded for the experimenter. Peripheral blood was sampled from the facial vein and analysed using a ScilVet Abc blood counter.

### Generation of HoxB8 progenitor cells

Cell lines were generated as described in^46^; in short, fresh bone marrow was cultivated in Optimem supplemented with recombinant mouse IL3 and IL6, spin-infected with *HoxB8*-encoding retrovirus and cultivated in RPMI-1640 complete (PFs) or Optimem complete (PNs) with either FLT3 (PF) or SCF (PN) supernatant. Primary cell culture was regularly checked for mycoplasma contamination with following primers: Forward: GGG AGC AAA CAG GAT TAG ATA CCC T; Reverse: TGC ACC ATC TGT CAC TCT GTT AAC CTC.

### Immunoblotting and Immunoprecipitation

Cells lysis, immunoblot analyses and immunoprecipitation were performed as previously described^68^; used antibodies are listed in Supplementary Material Table1.

### Flow cytometry and intracellular staining for phH3

Single-cell suspensions of the spleen, thymus and lymph node were achieved by smashing the whole organ through a 70 µm strainer (Corning, Cambridge, MA, USA, 352350) in PBS/2% FCS. Single-cell suspensions of the spleen were pelleted and resuspended in 1 ml red blood cell lysis buffer for erythrocyte depletion for 2 min at RT, washed with PBS, then stained with antibodies and quantified for cellularity using a hemocytometer (Neubauer) and trypan blue exclusion. Flow cytometric analysis of single-cell suspensions was performed on an LSR Fortessa (BD) and analysed using FlowJo® v10 software.

HoxB8 cells were fixed in 70 % ethanol and stored at −20°C. After two washes with PBS, cells were incubated for 15 min in PBS/0.25% Triton X-100 on ice for permeabilisation. Cells were washed in PBS/1% BSA and incubated for 60′ with anti-mouse phospho-Histone3 S10 mAb. Cells were washed and stained with ToPro-3 (100 nM) for DNA content analysis. List of Antibodies in Supplementary Material Table 3.

### Single-cell sequencing and data analysis

HoxB8-PN cell lines treated with Doxycycline were harvested at different time points and frozen in freezing media (FCS+10% DMSO) until further processing for whole-genome single-cell sequencing, performed on 24 cells/cell line using a NextSeq 500 (Illumina, San Diego, CA). Data analysis was performed as described previously^69^. Short, the sequencing data was aligned using the murine reference genome (GRCm38) along with the Bowtie2 (v2.2.4) software^70^. The AneuFinder (v1.10.1)^71^ was used for copy number variation analysis; libraries were processed, including GC correction, elimination of regions prone to artefacts, and validation of mappability. The divisive copy number calling algorithm was employed with a bin size of 1 Mb. The modal copy number state was determined based on the anticipated ploidy state. To calculate the aneuploidy score for each library, the average absolute deviation from the expected euploid copy number was computed per bin.

### Live cell imaging

For experiments shown in Fig S1 and S6, HoxB8 cells were seeded in media with or without 1 µg/ml Doxycycline and 1 µg/ml PI and imaged every 2h in an Incucyte S3 microscope using the 10x objective (Sartorius, Göttingen, Germany). The analysis function of the system was used to calculate the red object count (ROC), representing PI^+^ cells per mm², normalised to the area covered by the cells at start of imaging. Number of positive cells at the imaging start (t=0h) was subtracted from each time point.

For experiments shown in Figure 1G, H HoxB8-PF cells were seeded in media containing 1% methylcellulose in a 35mm petri-dish at a density of 400.000/ml and treated with or without 1 µg/ml Doxycycline and SPY650-DNA stain. Image acquisition was done with Zeiss Axio Observer microscope with 63x objective every 10 min in an environmental chamber set to 37°C. A total of 40 to 52 cells were randomly selected, and mitotic duration and cell fate were assessed manually using Fiji. The clear appearance of metaphase defined the onset of mitosis.

### Colony-forming assay

For colony-forming assays, MethoCult™ SF H4436 medium was used. Bone marrow was seeded with or without 1 µg/ml Doxycycline at a density of 10^3^ cells/ml/35 mm cell culture dish and incubated in the presence of serum-free medium with mouse cytokines: M-CSF for macrophages and G-CSF for granulocytes. MethoCult™ SF M3630 medium with IL-7 was used for B cell outgrowth. Colonies were counted 10-14 days post-seeding.

### Histology and immunohistochemistry

The intestine was transferred as a “swiss roll” to PBS/4% paraformaldehyde (PFA) for fixation and embedding. After 24h of fixation, tissue was transferred to 70% EtOH. Dehydration and processing were performed overnight via Shandon Citadel 1000 (50% EtOH, 70% EtOH, 80% EtOH, 2x 96%EtOH, 2x 100%EtOH, 2x Xylene), and then the tissue was embedded in paraffin. Histology blocks were cut sagittal with a rotation microtome (Leica Jung RM 2035), and sections were placed on silanised glass slides. Slides were dried for 30-60 min at 60°C. After deparaffinisation and rehydration of the slides, antigen retrieval was performed in 10mM NaCit-buffer, pH6 at 90°C. After washing with TBST, endogenous peroxidase activity was blocked with 3% H_2_O_2_ followed by permeabilisation and blocking of non-specific binding (5% goat serum in TBST) for 1h. 50 µl of the primary antibody (Supplementary Material Table 2) was added to the slides and incubated at 4°C overnight. Slides were washed with TBST, before incubation with the biotinylated secondary antibody (goat anti-rabbit 1:100; Dianova 111-065-144), for 2h at RT. Amplification was performed with the Vectastain ABC-Kit (Vector Laboratories, Cat# PK-6100) for 30min at RT followed by washing with TBST. Peroxidase substrate mix was prepared according to the ImmPACT kit (Vector Laboratories, Cat# SK-4105) manual, 100µl of the solution was added onto each slide. Counterstaining was done with Mayers Hematoxylin (Merck Cat# 109249). After that, slides were dehydrated and transferred to Xylene, then mounted with Histokitt. Images were taken with Zeiss Axioplan2 imaging at 10x and 20x magnification.

### Histopathological Examination

4μm sections of paraffin-embedded H&E stained intestine were examined for histological analysis by a board-certified pathologist.

| Colitis/Enteritis grade | Definition |
| --- | --- |
| 1 | increased apoptotic enterocytes and neutrophilic granulocytes in the *L. epithelialis* |
| 2 | Grade 1 plus: crypt abscesses (colon); increased inflammatory |
|  | infiltrate of the *L. propria* (ileum) |
| 3 | Grade 1 plus 2: erosions of the *L. epithelialis* |
| 4 | Grade 1-3 plus: ulceration |

Number of cells positive for phH3 or active Caspase-3 was assessed manually, with ImageJ, in at least 5-10 (colon) or 15-20 (small intestine) randomly chosen fields at 20x magnification per mouse. Images were acquired on a Zeiss Axioplan2 imaging microscope using ZENblue software. A mean is calculated for each mouse and genotype. All images were blinded before analysis.

### Statistical analysis

All statistical analyses were performed using Prism9 software (GraphPad, La Jolla, CA, USA). One-tailed paired Student *t-*test or analysis of variance (ANOVA) for multiple group comparisons. Data are shown as Mean ± SEM if not stated otherwise. Statistical significance is shown with symbols: * p-value <0.05, ** p-value <0.01, *** p-value 0.001, # p-value <0.0001

## Abbreviations

SAC: spindle assembly checkpoint
APC/C: Anaphase promoting complex
MCC: mitotic checkpoint complex
CIN: chromosomal instability
KO: knock out
OE: overexpression
Dox: Doxycycline
BM: bone marrow
SP: spleen
LN: lymph node
Thy: Thymus
HSPCs: Haematopoietic stem and progenitor cells

## Author contribution statement

GK performed experiments, analysed data, prepared figures, wrote the manuscript; FS & VB assisted with animal experiments; MD conducted histopathology assessment; RS provided MAD2 transgenic mice; DS & AT & FF conducted single-cell whole genome sequencing and analysed the results; SG established live-cell imaging experiments; AV designed research, analysed data, wrote the manuscript, conceived the study.

## Acknowledgements

We are grateful to I. Gaggl, C. Soratroi, J. Heppke, M. Fischer, N. Heinrich, M. Saurwein for excellent technical assistance or animal care; we also want thank to A. Strasser and L. Fava for sharing mice or reagents and all lab members for fruitful discussion. This work was supported by the Austrian Science Fund, FWF-funded project # I-3271 and the ERC AdG POLICE (787171).

